# Hepatic cholinergic T cell signaling mediates an adaptive response to alcohol-induced liver stress

**DOI:** 10.64898/2026.09.18.752097

**Authors:** Alexander J. Knights, Shanshan Liu, Evan J. Kim, Jun Wu

**Affiliations:** Life Sciences Institute, University of Michigan, Ann Arbor, MI, USA; Department of Orthopaedic Surgery, Washington University, St Louis, MO, USA; Center of Regenerative Medicine, Washington University, St Louis, MO, USA; Department of Molecular & Integrative Physiology, University of Michigan, Ann Arbor, MI, USA

## Abstract

Alcohol-associated liver disease (ALD) is a major cause of liver-related morbidity and mortality, yet no targeted, disease-modifying therapy exists and the immune mechanisms driving disease progression remain poorly defined. Acetylcholine-producing cholinergic immune cells have emerged as regulators of homeostasis across tissues and in disease, but their role in alcohol-induced hepatic stress is unknown. Here, we characterized the hepatic cholinergic immune landscape during alcohol-induced stress using flow cytometry and single-cell RNA-sequencing of ChAT-eGFP reporter mice. Alcohol feeding expanded hepatic cholinergic immune cells, with T cells constituting a substantial and dynamic fraction of this population that showed a broad increase in acetylcholine signaling and shifted towards a repressed, anergy-like transcriptional state. CD4 T cells underwent coordinated suppression of cholesterol biosynthesis, while CD8 T cells instead adopted a cytotoxic activation program. Together, these findings identify cholinergic T cell signaling, and cholesterol metabolism within CD4 T cells specifically, as a previously unrecognized adaptive response to alcohol-induced liver stress, nominating candidate therapeutic targets for a disease that is marred by critically limited treatment options.

## Introduction

Alcohol-associated liver disease (ALD) is a leading cause of chronic liver disease and liver-related mortality worldwide, and its global burden continues to rise in parallel with increasing rates of hazardous alcohol consumption(1). Despite this growing burden, therapeutic options remain profoundly limited: corticosteroids, the current standard of care for severe alcohol-associated hepatitis, provide only modest short-term survival benefit, and no targeted, disease-modifying therapy has been approved to date(2). A growing body of evidence implicates the hepatic immune compartment as a central driver of ALD pathogenesis. Kupffer cells and recruited monocyte-derived macrophages orchestrate the inflammatory, fibrotic, and metabolic derangements that characterize disease progression, and functional heterogeneity within these myeloid populations shapes clinical outcomes in ALD(3). Yet despite this increasing appreciation for immune contributions to ALD, how hepatic immune populations sense and adaptively respond to alcohol-induced stress, and what signals coordinate this response, remain incompletely understood.

One such coordinating signal is acetylcholine, classically considered a product of neuronal cholinergic transmission but now recognized as a mediator that immune cells can synthesize and release. Acetylcholine-producing, choline acetyltransferase (ChAT)-expressing immune populations have been identified in multiple tissues, where they act locally through nicotinic acetylcholine receptor signaling to shape tissue function independent of classical neural innervation. Cholinergic lymphocytes have been shown to regulate the response to viral infection(4), control vasodilation in the vasculature(5), and govern humoral immunity(6). In adipose tissue, our group and others have shown that non-neuronal cholinergic signaling regulates thermogenic and metabolic programs(7–16). We recently extended this paradigm to the liver, showing that cholinergic macrophages, including Kupffer cells, synthesize and release acetylcholine to activate nicotinic receptor signaling in neighboring hepatocytes, restraining lipogenic and inflammatory programs and protecting against the development of metabolic dysfunction-associated steatohepatitis(17). Furthermore, recent findings have uncovered a hepatic cholinergic immune network comprised of B cells, macrophages, and T cells, that governs liver regeneration(18). These findings establish the liver as a site of locally-produced, functionally consequential cholinergic immune signaling. However, whether this axis is engaged by alcohol-induced hepatic stress, and which cholinergic immune players are most strongly implicated in this response, has not been explored. Here, we characterize the cholinergic immune landscape of the liver during alcohol-induced stress, with a particular focus on T cells as a dynamic and responsive cholinergic immune population in this setting.

## Results

### Single-cell transcriptomic profiling of cholinergic cells in alcohol-induced liver stress

Flow cytometric profiling of liver non-parenchymal cells (NPCs) from ChAT-eGFP mice on an EtOH or pair-fed diet showed a statistically significant EtOH-induced increase in both the total abundance of ChAT-eGFP+ cells, and their proportion relative to all live NPCs (Figure 1, A and B). This result was corroborated using ChAT-eGFP mice crossed to ChAT-cre;RFP mice, with a shift towards more numerous RFP+ eGFP+ double positive cholinergic cells after an EtOH diet compared to pair-fed animals (Figure 1C). To understand the hepatic landscape of cholinergic cells at single-cell resolution, we then performed single-cell RNA-sequencing (scRNA-seq) on ChAT-eGFP+ NPCs from mice subjected to an EtOH diet or pair-fed controls. ChAT-eGFP+ NPCs were FACS-sorted, fixed, and prepared for multiplex scRNA-seq using the 10x GEM-X Flex platform (Figure 1D and Supplemental Figure 1A). Sequencing data was subjected to quality control and low-quality cells, debris, and doublets were filtered out to yield a total of 16,222 high-quality cells for analysis (Supplemental Figure 1B). Semi-supervised clustering of EtOH and pair-fed conditions combined yielded 10 cell clusters (0-9), identified by canonical marker gene expression and alignment with the ImmGen mouse immune cell RNA-seq database using the Cluster Identity Predictor (CIPR) tool(19) (Figure 1, E and F, and Supplemental Figure 1, C and D). Approximately 90% of cells registered non-zero expression of *Ptprc*, which encodes CD45, confirming our prior flow cytometry findings that the majority of ChAT-eGFP+ NPCs are of the hematopoietic/immune lineage(17) (Figure 1, G and H). Cell types identified included B cells, T cells (γδ and αβ), myeloid cells like macrophages and dendritic cells (DCs), as well as proliferating cells and red blood cells (RBCs) (Figure 1, E and F, and Supplemental Figure 1C). Comparing conditions, cholinergic signaling was significantly elevated in the EtOH group, calculated using an acetylcholine signaling gene module score (Figure 1I), and expression of *Chat*, the gene encoding choline acetyltransferase (ChAT), was significantly upregulated in the EtOH group (Figure 1J). Together, these data defined the major types of ChAT-eGFP+ cells in livers from EtOH and pair-fed mice, mostly represented by the hematopoietic lineage, and demonstrated elevated cholinergic signaling on an alcohol diet.

**Figure 1.**
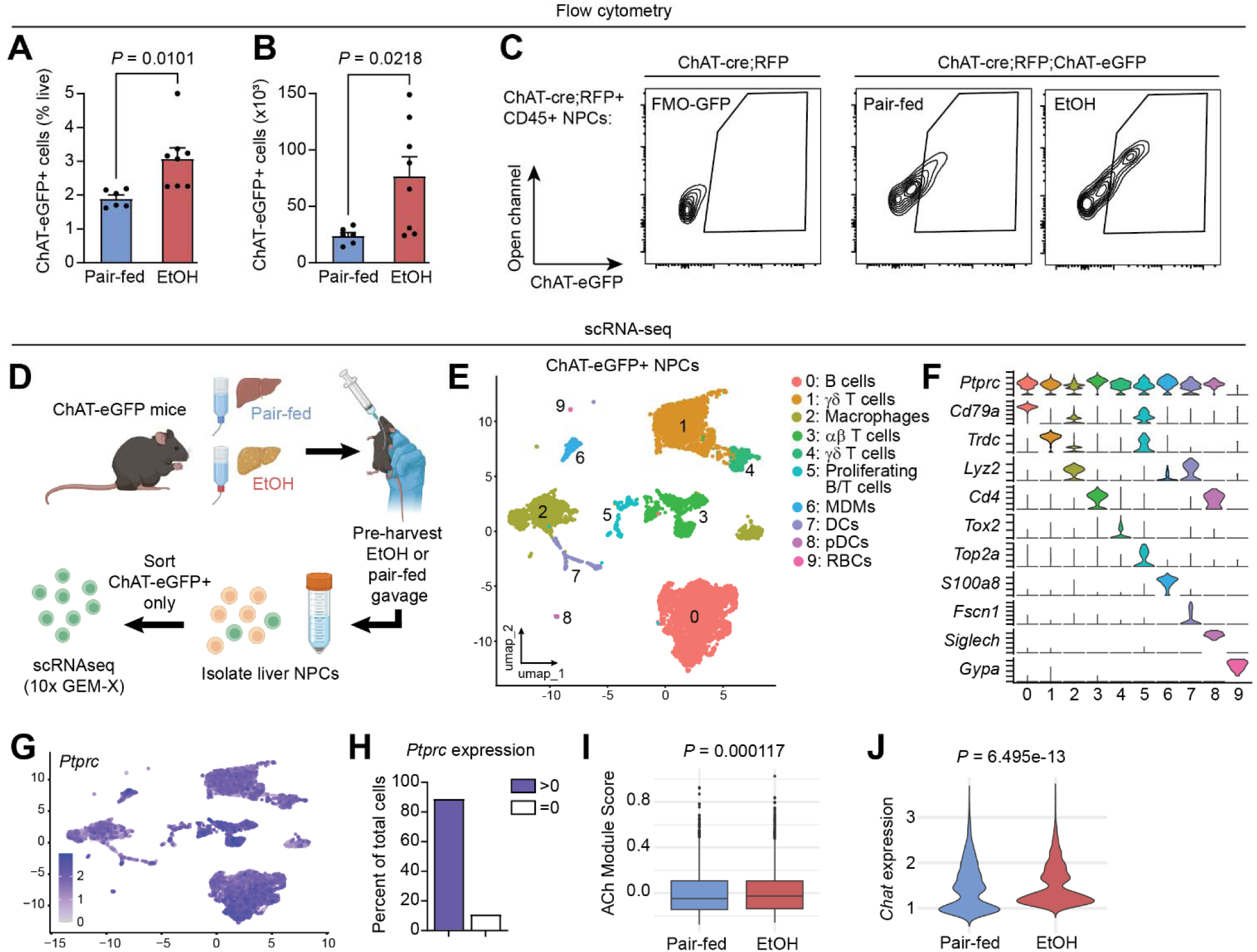
Single-cell transcriptomic profiling of cholinergic cells in alcohol-induced liver stress. (**A-B**) Flow cytometry analysis of ChAT-eGFP+ liver NPCs from EtOH or pair-fed mice (n=6-8), expressed as a percentage of total live cells (**A**) and total number of ChAT-eGFP+ cells (**B**). (**C**) ChAT-cre;RFP;ChAT-eGFP double reporter mice were subjected to an EtOH or pair-fed diet then the ChAT+ population assessed by flow cytometry. ChAT-cre;RFP+ CD45+ NPCs were gated on for ChAT-eGFP positivity, including a fluorescence-minus-one control for GFP (left). (**D**) Schematic showing the experimental design, resulting in scRNA-seq of sorted ChAT-eGFP+ liver NPCs from EtOH or pair-fed mice. (**E**) UMAP plot showing ChAT-eGFP+ NPC clustering and cell type naming of both conditions combined (16,222 total cells). (**F**) Canonical marker genes that define each cluster. (**G**) Expression of *Ptprc* (encoding CD45) in all cell clusters. (**H**) Percent of total cells expressing *Ptprc* at levels greater than zero, or equal to zero. (**I**) Composite expression score of an acetylcholine (Ach) gene module for all cells in the pair-fed or EtOH condition. Box-and-whisker plotting shows the median and interquartile range. (**J**) Violin plot of *Chat* (encoding choline acetyltransferase, ChAT) in all cells from pair-fed or EtOH conditions. For **A** and **B**, error bars represent mean±SEM and statistical testing was performed by two-tailed unpaired student’s t-tests. For **I** and **J**, statistical testing was performed by Wilcoxon rank sum test with continuity correction and exact *P* values are shown above each graph. MDMs: monocyte-derived macrophages; DCs: dendritic cells; pDCs: plasmacytoid dendritic cells; RBCs: red blood cells.

### Conservation of cholinergic immune cells in adipose tissue and liver

Given our previous transcriptomic profiling of ChAT-eGFP+ cells in the stromal vascular fraction (SVF) of murine inguinal white adipose tissue(10), and our new data from the liver, we sought to understand commonalities between the cholinergic cell landscape in both of these tissue settings. Data from our previous study of adipose SVF from mice housed at thermoneutrality(10) was integrated with the pair-fed liver scRNA-seq data from this study using Harmony, to assess ChAT-eGFP cells in these tissues. Semi-supervised clustering revealed four major cell types expressing ChAT-eGFP in adipose SVF and liver NPCs (Figure 2A). *Ptprc* (CD45) was broadly expressed across almost all cells, with stromal (*Pdgfra*, *Pdpn*) and endothelial (*Tie1*) markers confined to only a few cells across datasets (Figure 2B). Canonical markers were used to identify the major cell types, namely *Cd19* for B cells, *Cd3e* for T cells, *Lyz2* for myeloid cells, and *Eln* for stromal cells (Figure 2C). Cell types were broadly found in both tissues (Figure 2, D and E), with the most noticeable discrepancy in the ChAT-eGFP+ myeloid cell fraction, likely owing to the unique presence of Kupffer cells in the liver (which we previously identified as ChAT-eGFP+(17)), and their absence in adipose tissue. Lymphocytes comprised the majority of cholinergic cells in both adipose and liver with myeloid cells representing ~5-15% of the total ChAT-eGFP+ population. Of interest, cell cycle scoring (S and G2M phases) revealed the presence of proliferative ChAT-eGFP+ cells in both tissues, and across all four cell types (Figure 2, F-H). This integrated analysis demonstrated the shared cellular landscape of ChAT-eGFP+ cells in liver and adipose tissue, predominated by hematopoietic cells (T and B cells, myeloid cells), including cells with active proliferative status.

**Figure 2.**
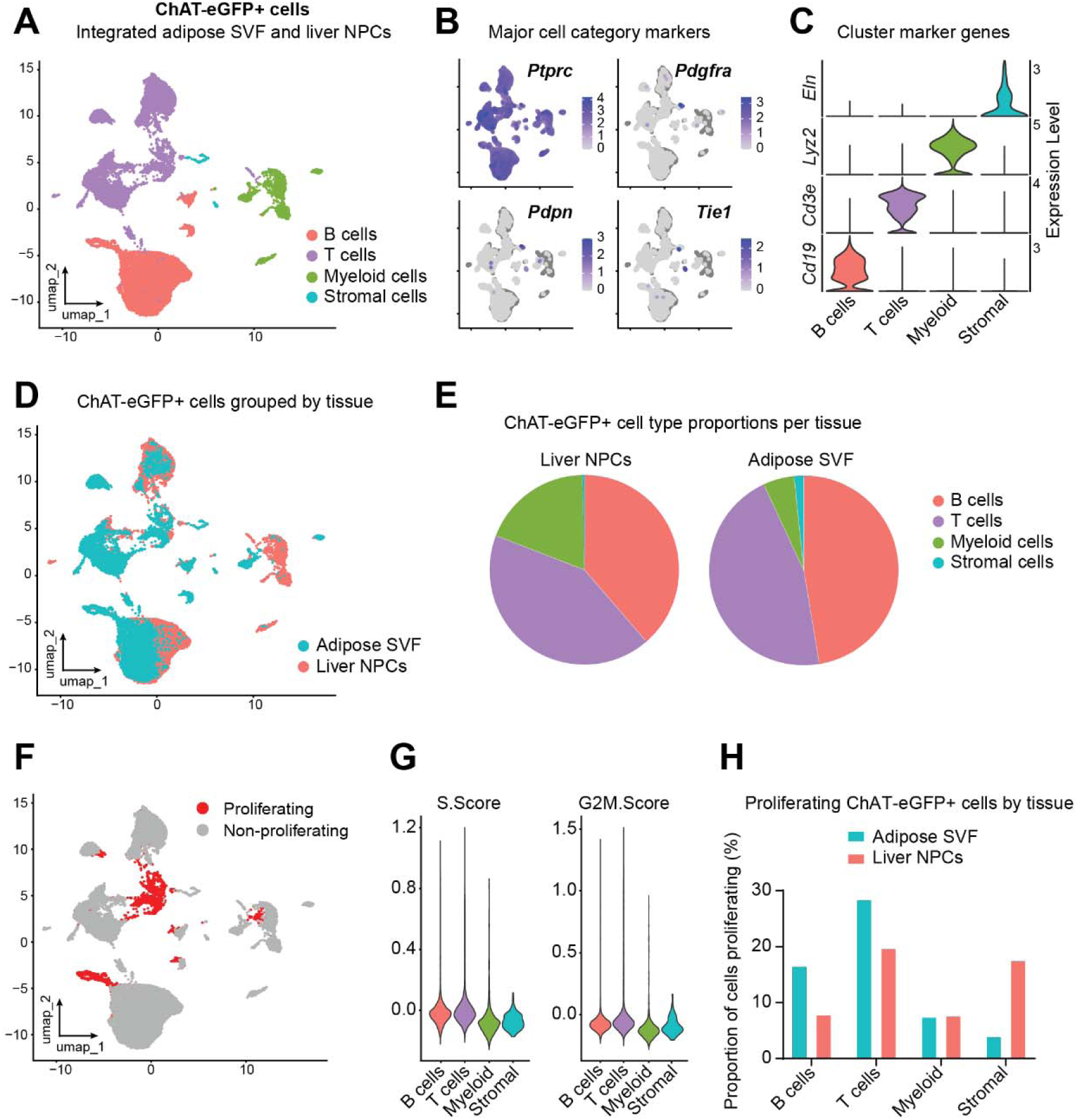
Conservation of cholinergic immune cells in adipose tissue and liver. (**A**) UMAP plot of integrated data from adipose stromal vascular fraction (SVF) and liver NPCs showing the major types of ChAT-eGFP+ cells. (**B**) Expression of genes that mark broad cell categories: *Ptprc* (immune/hematopoietic), *Pdgfra* and *Pdpn* (stromal), and *Tie1* (endothelial). (**C**) Expression of marker genes that define each cluster from (A). (**D**) UMAP plot showing the distribution of ChAT-eGFP+ cell clusters grouped by tissue source. (**E**) Proportional breakdown of major ChAT-eGFP+ cell types by tissue source. (**F**) UMAP plot showing cells classified as proliferating (S or G2M phase). (**G**) Cell cycle module scores by cell type. (**H**) Proportion of each ChAT-eGFP+ cell type classified as proliferating, per tissue source.

### Cholinergic T cell perturbation in alcohol-induced liver stress

Based on our scRNA-seq data, T cells comprise approximately half of the cellular pool of ChAT-eGFP+ NPCs in the liver (Figure 2E), yet nothing is known about their pathophysiological functions in this setting. Flow cytometry using ChAT-eGFP mice showed a statistically significant increase in ChAT-eGFP+ T cells following an EtOH diet (Figure 3A). Using our scRNA-seq data to focus on the cholinergic T cell fraction and remove transcriptional noise introduced by other cell types, we computationally subset the T cell clusters (1, 3, 4), marked by expression of *Cd3e* (Figure 3B). Cluster 5 (from Figure 1E) was not included given its mixed signature of proliferating B and T cells (Supplemental Figure 2A). Directly comparing ChAT-eGFP+ T cells from the EtOH condition with those from the pair-fed condition, we found 889 differentially-expressed genes (DEGs; padj<0.05 and Log2FC>|0.585|) (Figure 3C and Supplemental Figure 2B). Scoring of a composite acetylcholine signaling gene module, which includes *Chat*, showed significantly elevated activity in T cells from the EtOH group, consistent with an adaptive disease-induced response (Figure 3D). Pathway analysis revealed a broad T cell anergy-like state in the EtOH group, with repressed activation, complement activity, and biosynthesis (Supplemental Figure 2, C and D). Apoptotic pathways were also overrepresented in the EtOH condition. Proliferating, CD4, CD8, and γδ ChAT-eGFP+ T cell subsets were identified, with a marked distinction between TCRαβ and TCRγδ populations (Figure 3, E-G). Of these ChAT-eGFP+ subsets, CD4 T cells showed the greatest proportional increase on the EtOH diet (Figure 3H and Supplemental Figure 2E), and thus we decided to further interrogate this population.

**Figure 3.**
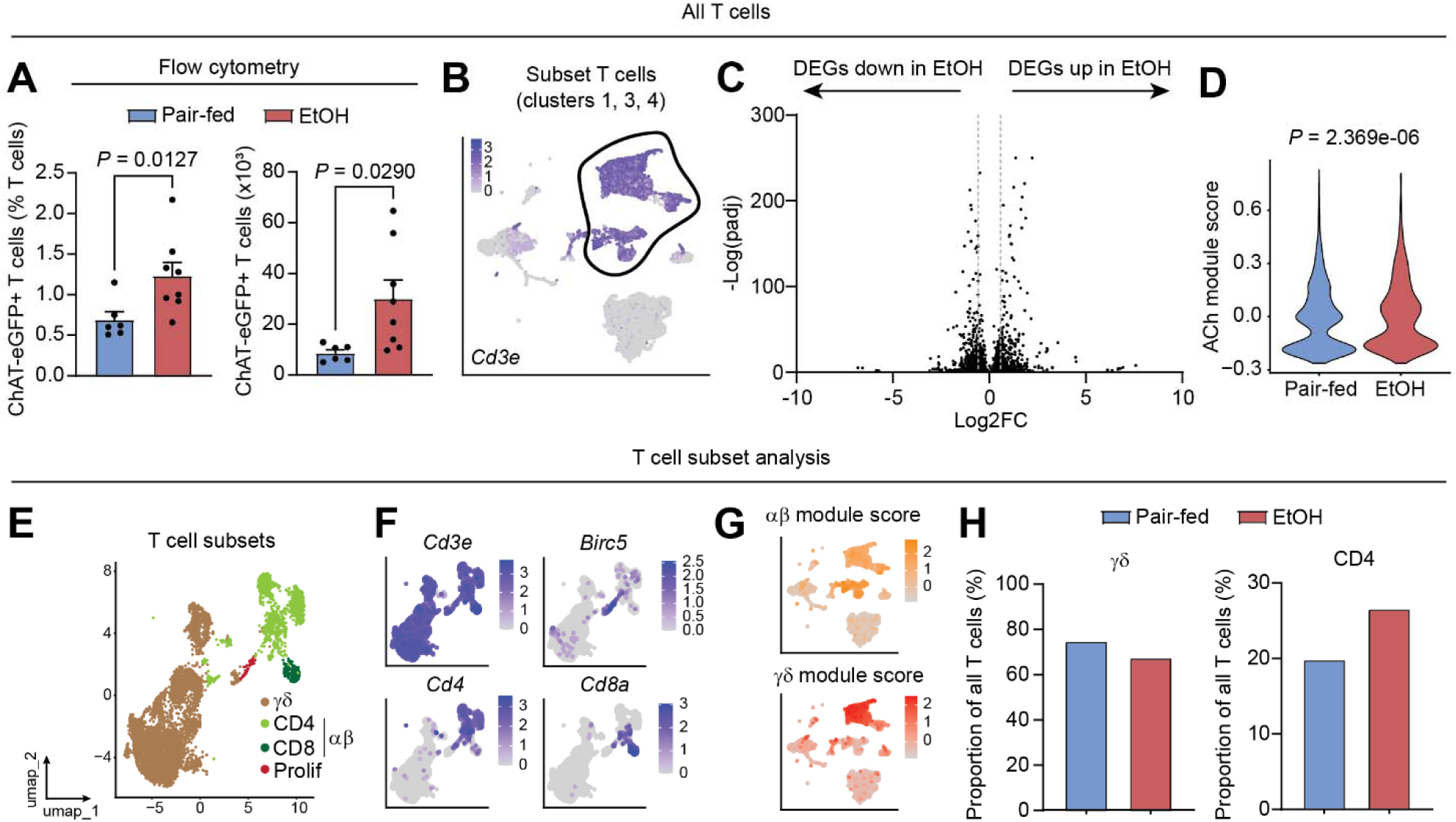
Cholinergic T cell perturbation in alcohol-induced liver stress. (**A**) Flow cytometry analysis of ChAT-eGFP+ liver T cells from mice fed an EtOH or pair-fed diet (n=6-8), expressed as a percentage of total T cells (left) and absolute number of ChAT-eGFP+ T cells (right). (**B**) ChAT-eGFP+ T cell clusters (1, 3, 4) were computationally subset based on expression of *Cd3e*. (**C**) Volcano plot showing DEGs derived from pseudobulk differential expression analysis of all T cells in the EtOH versus the pair-fed condition. Positive Log2FC indicates genes that were upregulated in EtOH. Vertical dashed lines intersect the x axis at Log2FC −0.585 and +0.585. (**D**) Composite expression score of an acetylcholine (Ach) gene module for all T cells in the pair-fed or EtOH condition. (**E**) ChAT-eGFP+ T cells divided into four clusters comprised of γδ T cells, αβ T cells (CD4 and CD8), and proliferating T cells (Prolif). (**F**) Feature plots showing expression of *Cd3e, Cd4, Cd8*, and *Birc5* in T cell subsets. (**G**) Feature plots of module scores for γδ- or αβ-associated genes, overlaid onto the UMAP of all ChAT-eGFP+ NPCs. (**H**) Proportional abundance change for γδ (left) and CD4 (right) ChAT-eGFP+ T cell subsets in the EtOH condition versus pair-fed. For **A**, error bars represent mean±SEM and statistical testing was performed by a Mann-Whitney test (left) or a two-tailed unpaired student’s t-test (right). For **D**, statistical testing was performed by Wilcoxon rank sum test with continuity correction, with exact *P* value shown.

ChAT-eGFP+ CD4 T cells had an elevated acetylcholine module score after EtOH diet and 692 statistically significant DEGs were detected between conditions (padj<0.05 and Log2FC>|0.585|) (Figure 4, A-C, and Supplemental Figure 2B). Gene ontology pathway analyses revealed suppression of cholesterol biosynthesis as a key recurring theme in the EtOH condition, along with mitochondrial stress, autophagy, and apoptosis (Figure 4D and Supplemental Figure 3, A and B). A gene module for cholesterol biosynthesis was curated and showed significant upregulation in EtOH, with genes encoding cholesterol synthetic enzymes overwhelmingly downregulated in this condition (Figure 4, E and F). This is salient given the widely reported crucial role of cholesterol for T cell activation, signaling, and motility(20). Gene set enrichment analysis (GSEA) similarly identified cholesterol homeostasis as negatively enriched in CD4 T cells from the EtOH group (Supplemental Figure 3B). Analysis of leading-edge genes driving this negative enrichment confirmed their global downregulation, with the notable and potent downregulation of *Srebf2* (Figure 4G), a transcription factor that controls cholesterol biosynthesis, providing a putative mechanism for the observed consistent suppression of cholesterol biosynthesis in ChAT-eGFP+ CD4 T cells after EtOH diet. ChAT-eGFP+ CD8 T cells, on the other hand, showed no apparent deregulation of cholesterol homeostasis nor adaptive elevation of acetylcholine signaling, but instead showed a robust cytotoxic activation program (Supplemental Figure 3, C-H). While overall, ChAT-eGFP+ T cells showed a broad suppression of activation and function following EtOH diet, discrimination between cellular subsets provided greater resolution and insight into the dysregulated functions induced in alcohol-associated liver disease, including notable impairment of cholesterol homeostasis in the CD4 population.

**Figure 4.**
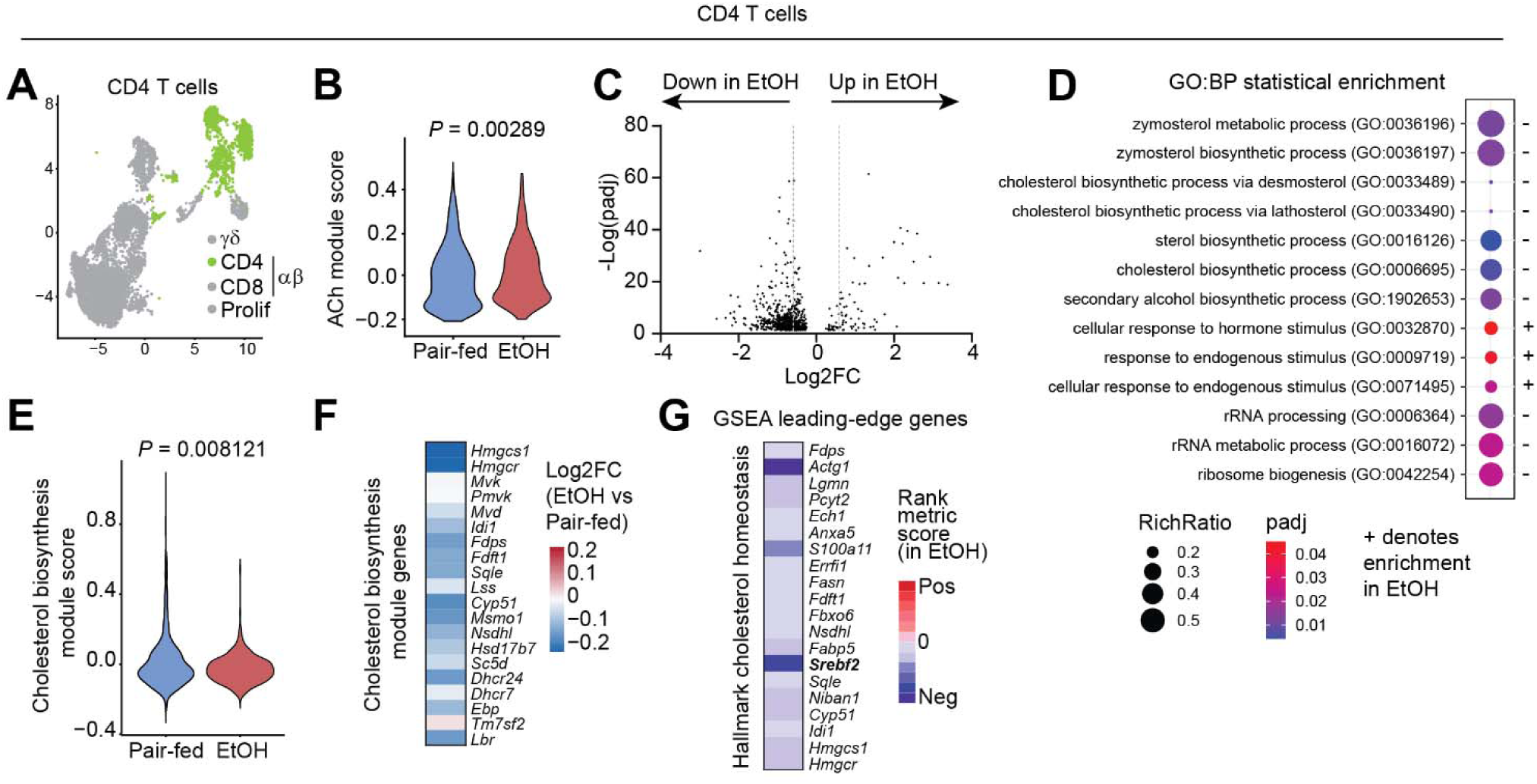
Cholinergic CD4 T cells are dysregulated in alcohol-induced liver stress. (**A**) UMAP showing CD4 T cells colored green. (**B**) Composite expression score of an acetylcholine (Ach) gene module for CD4 T cells in the pair-fed or EtOH condition. (**C**) Volcano plot showing DEGs derived from differential expression analysis of all CD4 T cells in the EtOH versus the pair-fed condition. Positive Log2FC indicates genes that were upregulated in EtOH. Vertical dashed lines intersect the x axis at Log2FC −0.585 and +0.585. (**D**) Bubble plot of significantly enriched biological pathways derived from statistical enrichment analysis of CD4 T cell DEGs (padj<0.05) between conditions. The Gene Ontology: Biological Processes (GO:BP) annotation set was used. The plus (+) symbol denotes terms that were enriched in the EtOH condition, and size and color of each bubble represent RichRatio and significance of each term, respectively. (**E**) Composite expression score of a manually curated Cholesterol biosynthesis gene module for CD4 T cells in the pair-fed or EtOH condition. (**F**) Heatmap showing fold enrichment (Log2FC) of each gene comprising the Cholesterol biosynthesis module, in CD4 T cells from EtOH compared to pair-fed. (**G**) Heatmap showing the rank metric score (in the EtOH condition) for significant leading-edge genes derived from gene set enrichment analysis (GSEA) of the Hallmark cholesterol homeostasis gene set. For **B** and **E**, statistical testing was performed by Wilcoxon rank sum test with continuity correction and exact *P* values are shown above each graph.

## Discussion

In this study, we identified cholinergic immune cells in the liver that expand in response to alcohol-induced hepatic stress, and we showed that T cells constitute a substantial and dynamic fraction of this population. Using single-cell transcriptomic profiling, we found that alcohol feeding induced a broad increase in acetylcholine signaling across ChAT-eGFP+ T cells, corresponding to an anergy-like transcriptional state marked by suppressed activation, complement, and biosynthetic programs. Within this population, CD4 and CD8 T cells diverged sharply: CD4 T cells underwent a pronounced, coordinated suppression of cholesterol biosynthesis, driven in part by downregulation of the master transcriptional regulator *Srebf2*, whereas CD8 T cells instead adopted a robust cytotoxic activation program. We further showed that the cholinergic immune landscape is broadly conserved between liver and adipose tissue, with lymphocytes representing the predominant non-neuronal ChAT-eGFP+ population in both tissues. These data support a model in which cholinergic T cell signaling constitutes an adaptive, disease-restraining response to alcohol-induced hepatic stress, with cholesterol metabolism emerging as a key regulated node specifically within the CD4 compartment. Further investigation aims to uncover whether this is indeed a protective response, and if this response becomes exhausted or aberrant over time.

The parallels we observe between cholinergic immune populations in the liver and adipose tissue add to an emerging view that acetylcholine-producing immune cells act as a conserved regulatory node linking metabolic tissues to local immune and stress responses. Cholinergic signaling from immune-derived acetylcholine has been implicated in shaping metabolic and inflammatory tone across multiple organs, suggesting that this axis may represent a broader mechanism by which the immune system calibrates tissue homeostasis under metabolic challenge, even as the specific molecular consequences of this signaling appear to vary by tissue and cell type.

These findings carry particular significance given the near-total absence of targeted therapies for ALD. By identifying cholinergic T cell signaling, and the regulation of cholesterol metabolism within this population, as a previously unrecognized adaptive axis in alcohol-induced liver stress, our findings reveal new candidate pathways for therapeutic investigation in an area where treatment options remain critically limited.

## Methods

### Sex as a biological variable

For flow cytometry experiments, both male and female mice were used, however sample sizes were not powered to disaggregate sex-dependent effects. For scRNA-seq, only male mice were used.

### Statistics

For flow cytometry data, the sample size is indicated in the figure legend, where n represents the number of mice used. Normality testing was performed (Shapiro-Wilk test) prior to unpaired Mann-Whitney or student t-tests, as indicated in Figure legends corresponding to each panel. Error bars represent the mean ± the standard error of the mean (SEM). For scRNA-seq data, Wilcoxon rank sum testing with continuity correction was performed to compare gene expression or gene module expression in pair-fed versus EtOH. Statistical enrichment and statistical overrepresentation testing of pathways used Fisher’s exact tests with false discovery rate correction to generate adjusted *P* values (*Padj*). For gene set enrichment analysis (GSEA), a weighted Kolmogorov-Smirnov statistic was used to generate a final enrichment score and *P* value that was corrected using false discovery rate, yielding an adjusted *P* value (termed q-val). In all instances, exact *P* or *Padj* values are given, with *P* (or *Padj*) < 0.05 considered statistically significant unless otherwise noted.

### Study approval

All animal work was performed in accordance with protocols approved by the University of Michigan Institutional Animal Care and Use Committee (IACUC).

### Mice

Wild-type mice used in this study were C57BL/6J mice (#000664 from Jackson Laboratories), and Ai14 (#007914), Chat-Cre (#031661), ChAT^BAC^-eGFP (#007902) mice were also used. All animals were housed under a 12/12 h light/dark cycle at ambient temperature (23°C) or in an environmental chamber at 30°C (thermoneutrality), where indicated. Age-matched mice were fed a Lieber-DeCarli liquid diet (Bio-Serv, F1258SP, with 5% EtOH) or were pair-fed a control diet (Lieber-DeCarli liquid diet supplemented with isocaloric maltose, Bio-Serv, F1259SP) after gradual acclimation. In accordance with the protocol outlined by the National Institute on Alcohol Abuse and Alcoholism (NIAAA), as described previously(21), mice were fed with 5% EtOH-containing Lieber-DeCarli liquid diet or pair-fed the liquid diet for 10 days. On the final day, mice were administrated with 5 g/kg body weight EtOH or isocaloric maltose solution by oral gavage and harvested 4 h later. Following the gavage, mice were housed at thermoneutrality, prior to harvest.

### Flow cytometry and cell sorting

Livers were harvested from ChAT-eGFP mice after and EtOH or pair-fed diet regimen, as above. Liver digestion and hepatocyte removal to yield NPCs was performed as per our previously reported protocol(17). NPCs were stained with a cocktail of antibodies comprised of CD45-BV650 (clone 30-F11), CD11b-BV605 (clone M1/70), CD19-PECy7 (clone 6D5), F4/80-APC/R700 (clone BM8), and CD3-APCFire750 (clone 17A2), alongside TOPRO3 as a live/dead dye. NPCs from wild-type mice were used for unstained, single-stained, and fluorescence-minus-one controls, which were used to inform gating. Flow cytometry was performed on a BD Fortessa and analysis performed using FlowJo v10.

### Single-cell RNA-sequencing (scRNA-seq)

Livers were harvested from male ChAT-eGFP mice after an EtOH or pair-fed diet regimen, as above. Five mice were pooled into a single sample per condition. Following tissue digestion and removal of hepatocytes, NPCs were subjected to cell sorting in FACS buffer containing 3 µM DAPI to discriminate live from dead cells. NPCs from wild-type mice were used for unstained and DAPI-only control tubes, to set the ChAT-eGFP+ gate. A ThermoFisher Bigfoot sorter was used with a 100 µm nozzle. A total of 50,000 ChAT-eGFP+ NPCs from each condition were sorted directly into 500 µL of Fixation Buffer B from the 10x Genomics GEM-X Flex Sample Preparation v2 Kit (PN-1000781) then fixed for 24 h at 4°C. After 24 h, 0.5 mL of Additive C was added, followed by Quench Buffer B. A 0.1x volume of Enhancer was then added, along with glycerol to a final concentration of 10%. Cell samples were then given to the University of Michigan Advanced Genomics Core for counting, library preparation, and sequencing. 10,000 target cells were input per condition and sequencing was performed on an Illumina NovaSeq X Plus at 28x90bp.

### scRNA-seq analysis

#### Liver NPC scRNA-seq quality control and analysis

Sequencing data for the pair-fed and EtOH samples were first processed in 10x CellRanger which showed that 100% of initial cell barcodes passed high occupancy GEM filtering and 92.8% of reads were confidently mapped in cells. The R package Seurat v5(22) was used for bioinformatic analysis of the dataset, and filtered count matrices for both sample conditions were read into Seurat using the function *Read10X* followed by creation of the Seurat object using *CreateSeuratObject* (min.features = 100, min.cells = 10). Violin plots for nFeatures, nCount, and mito.ratio (percent.mito) were generated using the *VlnPlot* function prior to filtering, then thresholds were defined as nFeature_low = 200; nFeature_high_perc = 0.93 (i.e. cells above the 93rd percentile of nFeatures were excluded); nCount_low = 100; nCount_high = 15000; mitoRatio_high = 100 (i.e. no mito.ratio cutoff was used). Samples were merged and the filtered Seurat object was then subjected to a parameter sweep across a range of dimensionality (20, 30, and 40 principal components) and clustering resolutions (0.1, 0.25, 0.5, 0.75, and 1.0) using Seurat’s *RunUMAP*, *FindNeighbors*, and *FindClusters* functions. UMAP plots were generated and visually inspected for each dimension/resolution combination to guide selection of final clustering parameters. A combination of targeted feature plots of cell type marker genes (using *FeaturePlot*) and genes unique to each cluster (from *FindAllMarkers*) were used to determine cluster identities based on semi-supervised clustering, with final settings of 40 dimensions and 0.1 resolution chosen, resulting in 10 clusters (0-9). The Cluster Identity Predictor (CIPR) tool(19) was used to cross-validate our cluster annotations based on gene signatures of pre-sorted murine cell types from the ImmGen database.

To enable direct visual comparison of UMAP distributions between conditions, cells were randomly downsampled in Seurat (*WhichCells* with downsample; seed = 42) to the smaller of the two condition-level cell counts, yielding equal numbers of cells per condition for visualization. *Ptprc* expression (encoding CD45) was visualized on the UMAP embedding, and the percentage of cells with detectable (raw count > 0) *Ptprc* expression was calculated. *Chat* expression was compared between conditions among *Chat*-expressing cells only (SCT-normalized expression > 0), and a composite acetylcholine signaling gene module score was calculated across conditions using Seurat’s *AddModuleScore* function (using a curated gene list from our previous publication(9): *Chat, Ache, Bche, Slc44a4, Slc18a3, Slc5a7, Tacr2, Adora2a, Chdh, Chka*). Assessment of statistical significance in gene or module expression was performed using the Wilcoxon rank-sum test.

For analysis of ChAT-eGFP+ T cells, clusters 1, 3, and 4 were computationally subset to remove variance derived from non-T cell clusters. T cell clusters were defined by *Cd3e* expression and gene signature derived from *FindAllMarkers*; the proliferating B/T cell cluster was not included due to an overwhelming cell cycling signature and B cell contamination. Pseudobulk differential expression analysis of T cells between pair-fed and EtOH conditions was performed using the *FindMarkers* function with FDR correction, yielding differentially expressed genes (DEGs) with an adjusted p-value (*padj*). For genes where *padj* = 0, the −log(*padj*) was artificially capped at 250 for visualization via volcano plot. Resulting DEG lists were used for downstream pathway enrichment analysis.

Within the subsetted T cell object, subclustering identified major subtypes of T cells. First, the *AddModuleScore* function was used to determine whether clusters were αβ or γδ T cells based on expression of *Trac*, *Trbc1*, *Trbc2*, *Trdc*, *Trgc1*, and *Trgc2*. Further, expression of targeted marker genes (*Birc5*, *Cd8a*, *Cd3e, Cd4*) was used to identify proliferating, CD4, CD8, and γδ clusters, and the number of cells in each subcluster for each condition was enumerated. *FindMarkers* was also used to assess differential gene expression within the CD4 and CD8 subclusters, and the resulting DEG lists were used for downstream pathway enrichment analysis.

PantherDB(23) was used for performing pathway enrichment analysis using the *Mus musculus* background gene list. Statistical enrichment testing of Gene Ontology (Biological Processes) used an input of all genes (ranked by log2FC and *padj*) from differential expression analysis. Bubble plots of enriched pathway terms were generated using the ggplot2 R package, where color scale denotes *padj* and bubble size represents RichRatio, which is the number of significant genes from the DEG list found in the term divided by the total number of genes in the pathway term. Statistical overrepresentation testing using the Fisher test was performed using input gene lists containing DEGS with *padj*<0.05 and log2FC>|0.585|. Bubble plots were made with ggplot2, where color scale represents *padj* of each term, and bubble size represents Fold Enrichment, which is the magnitude of the effect size for any given term (how many times more frequently genes from a specific category appear in your input DEG list compared to what you would expect purely by random chance). For both statistical enrichment and overrepresentation testing in PantherDB, false discovery rate correction was applied, with only terms showing *padj* (FDR) < 0.05 shown.

The *AddModuleScore* function was used to compare expression of a cholesterol gene module comprised of the following genes: *Hmgcs1, Hmgcr, Mvk, Pmvk, Mvd, Idi1, Fdps, Fdft1, Sqle, Lss, Cyp51, Msmo1, Nsdhl, Hsd17b7, Sc5d, Dhcr24, Dhcr7, Ebp, Tm7sf2, Lbr*. A Wilcoxon rank-sum test was used to assess statistical significance.

For gene set enrichment analysis, the GSEA desktop app (v4.4.0)(24) was used with the mouse MSigDB database and Hallmark gene sets. A .rnk file of all genes from differential expression analysis was used as input, where rank was defined as gene name and sign(log2FC) * −log10(pval). The GSEA preranked analysis was run using the default settings. Gene set enrichment was defined as q-value (FDR) < 0.25, and positively and negatively enriched significant gene sets were plotted based on their normalized enrichment score (NES) with bubble plots, where color scale represents q-val (FDR) and bubble size represents the number of genes in each set that were found in the input list. Leading edge genes in the Hallmark Cholesterol Homeostasis pathway were analyzed using the GSEA in-built leading edge gene analysis tool, and a heatmap generated based on rank metric score.

#### Integrated analysis of adipose SVF and liver NPCs

To study the ChAT-eGFP+ cell landscape in steady-state liver and adipose tissue, we integrated the scRNA-seq data from the pair-fed liver NPC condition of this study with scRNA-seq data from adipose SVF of ChAT-eGFP mice housed at thermoneutrality that we have previously published(10) (GSE303687). For both datasets, filtered count matrices were imported into Seurat using *Read10X* and *CreateSeuratObject* (min.features = 100). Cell barcodes were renamed with sample-specific prefixes using *RenameCells* to ensure uniqueness across samples, and metadata columns were added or standardized to track sample ID and tissue origin. For each sample, the percentage of mitochondrial transcripts (mito.ratio) was calculated per cell using *PercentageFeatureSet*. Quality control metrics (nFeature_RNA, nCount_RNA, and mito.ratio) were visualized using violin plots. Sample-specific filtering thresholds were applied as follows: liver samples retained cells with 200 < nFeature_RNA < 6000 and mito.ratio < 15%, and SVF samples retained cells with 200 < nFeature_RNA < 5000 and mito.ratio < 7.5%. To reduce object size prior to integration, filtered objects were slimmed using the *DietSeurat* function to retain the RNA assay counts while removing stored data layers, scaled data, and dimensionality reductions. Filtered samples were merged into a single Seurat object, which was then normalized via the *NormalizeData* function, highly variable features were identified (*FindVariableFeatures;* 3000 features), then data were scaled using variable features only (*ScaleData*), followed by principal component analysis (*RunPCA*; 50 PCs). Batch effects were corrected and datasets were integrated using the R package Harmony(25) with “sample” specified as the grouping variable, applied to PCs 1–30. UMAP embeddings and a shared nearest neighbor graph were computed in Harmony space (*RunUMAP* and *FindNeighbors*; dims = 1–30). Graph-based clustering was performed using *FindClusters* across multiple resolutions, with cluster assignments saved for comparison across resolutions. UMAPs were visualized colored by cluster number, tissue, and condition, then *FindAllMarkers* was used in combination with targeted feature plots (*FeaturePlot*) of marker genes to determine the identities of each cluster. Clusters in the integrated Seurat object were pseudoclustered to represent the four major ChAT-eGFP+ cell types: T cells, B cells, myeloid cells, and stromal cells. Stacked violin plots of cell type markers were generated using the *VlnPlot* function, and identification of proliferating cells was performed using the *CellCycleScoring* function based on curated mouse lists of S phase genes (*Mcm4, Rrm2, Mrpl36, Chaf1b, Gmnn, Exo1, Msh2, Cdc45, Uhrf1, Fen1, Rad51ap1, Cdc6, Cenpu, Hells, Dscc1, Slbp, Ubr7, Gins2, Dtl, Nasp, Wdr76, Tipin, Cdca7, Blm, Pola1, Mcm6, Usp1, Mcm5, Ung, Prim1, Clspn, Tyms, Polr1b, Rad51, Pcna, Casp8ap2, Mcm7, Rrm1, Ccne2, Rfc2, E2f8*) and G_2_/M phase genes (*Cbx5, Cdc25c, Gtse1, Smc4, Dlgap5, Ctcf, Cdca2, Kif20b, Cdca3, Tacc3, Hmgb2, Cks2, Ckap2, Top2a, G2e3, Ncapd2, Ttk, Cdk1, Gas2l3, Anp32e, Psrc1, Cenpa, Cks1b, Nek2, Birc5, Lbr, Kif23, Kif11, Aurkb, Rangap1, Cenpe, Hjurp, Cdca8, Ccnb2, Ndc80, Tpx2, Hmmr, Cenpf, Cdc20, Ect2, Anln, Kif2c, Mki67, Ube2c, Tubb4b, Pimreg, Nusap1, Nuf2, Aurka, Bub1, Jpt1, Ckap5, Ckap2l*).

### Data availability

The adipose SVF scRNA-seq dataset is available at the NCBI Gene Expression Omnibus (GEO) under the accession GSE303687. The liver NPC scRNA-seq dataset will be deposited to GEO following peer review.

## Author contributions

Designed research studies: AJK, JW

Conducted experiments: AJK, SL

Acquired data: AJK

Analyzed data: AJK, EJK

Wrote the manuscript: AJK, JW

## Acknowledgements

We are grateful to the University of Michigan Flow Cytometry Core for access to their equipment and services.

## References

1. Danpanichkul P. The global epidemiology of alcohol-associated liver disease. Hepatol Commun. 2026;10(5).

2. Rattan P. Current and emerging therapies for acute alcohol-associated hepatitis. Aliment Pharmacol Ther. 2022;56(1):28–40.

3. Sasaki K. Kupffer cell diversity maintains liver function in alcohol-associated liver disease. Hepatology. 2025;81(3):870–87.

4. Cox MA. Choline acetyltransferase-expressing T cells are required to control chronic viral infection. Science. 2019;363(6427):639–44.

5. Tarnawski L. Cholinergic regulation of vascular endothelial function by human ChAT(+) T cells. Proc Natl Acad Sci U S A. 2023;120(14):e2212476120.

6. Nechanitzky D. Lymphocyte-derived cholinergic circuits modulate germinal center output and B cell activation. Nat Immunol. 2026;27(4):854–66.

7. Jun H. An immune-beige adipocyte communication via nicotinic acetylcholine receptor signaling. Nat Med. 2018;24(6):814–22.

8. Jun H. Adrenergic-Independent Signaling via CHRNA2 Regulates Beige Fat Activation. Dev Cell. 2020.

9. Knights AJ. Acetylcholine-synthesizing macrophages in subcutaneous fat are regulated by beta2 - adrenergic signaling. EMBO J. 2021;40(24):e106061.

10. Knights AJ. Transcriptomic plasticity of cholinergic adipose macrophages in the acute thermogenic response. J Biol Chem. 2025;301(12).

11. Ma Y. Immune cell cholinergic signaling in adipose thermoregulation and immunometabolism. Trends Immunol. 2022;43(9):718–27.

12. Ma Y. CHRNA2: a new paradigm in beige thermoregulation and metabolism. Trends Cell Biol. 2022;32(6):479–89.

13. Knuth CM. Subcutaneous white adipose tissue independently regulates burn-induced hypermetabolism via immune-adipose crosstalk. Cell Rep. 2024;43(1):113584.

14. Zhu K. Chrna2-driven CRE Is Expressed in Beige Adipocytes. Endocrinology. 2024;166(1).

15. Liu S. Nicotinic Acetylcholine Receptor Signaling Activates Beige Adipocytes and Mediates Systemic Metabolism. Diabetes. 2026;75(9):1511–24.

16. Meng W. The miR-182-5p/FGF21/acetylcholine axis mediates the crosstalk between adipocytes and macrophages to promote beige fat thermogenesis. JCI Insight. 2021;6(17).

17. Jun H. Signaling through the nicotinic acetylcholine receptor in the liver protects against the development of metabolic dysfunction-associated steatohepatitis. PLoS Biol. 2024;22(7):e3002728.

18. Modares NF. B cell-derived acetylcholine promotes liver regeneration by regulating Kupffer cell and hepatic CD8(+) T cell function. Immunity. 2025;58(5):1201–16 e7.

19. Ekiz HA. CIPR: a web-based R/shiny app and R package to annotate cell clusters in single cell RNA sequencing experiments. BMC Bioinform. 2020;21(1):191.

20. Bietz A. Cholesterol Metabolism in T Cells. Front Immunol. 2017;8:1664.

21. Bertola A. Mouse model of chronic and binge ethanol feeding (the NIAAA model). Nat Protoc. 2013;8(3):627–37.

22. Hao Y. Dictionary learning for integrative, multimodal and scalable single-cell analysis. Nat Biotechnol. 2024;42(2):293–304.

23. Mi H. Large-scale gene function analysis with the PANTHER classification system. Nat Protoc. 2013;8(8):1551–66.

24. Subramanian A. Gene set enrichment analysis: a knowledge-based approach for interpreting genome-wide expression profiles. Proc Natl Acad Sci U S A. 2005;102(43):15545–50.

25. Korsunsky I. Fast, sensitive and accurate integration of single-cell data with Harmony. Nat Methods. 2019;16(12):1289–96.

