## Supplementary Materials for "Hepatic cholinergic T cell signaling mediates an adaptive response to alcohol-induced liver stress"

**Supplemental material**

Knights *et al*, 2026

Supplemental Figure 1: scRNA-seq of ChAT-eGFP+ NPCs.

Supplemental Figure 2: Impaired function of cholinergic T cells in liver stress.

Supplemental Figure 3: Cholinergic CD4 and CD8 T cell responses in liver stress.

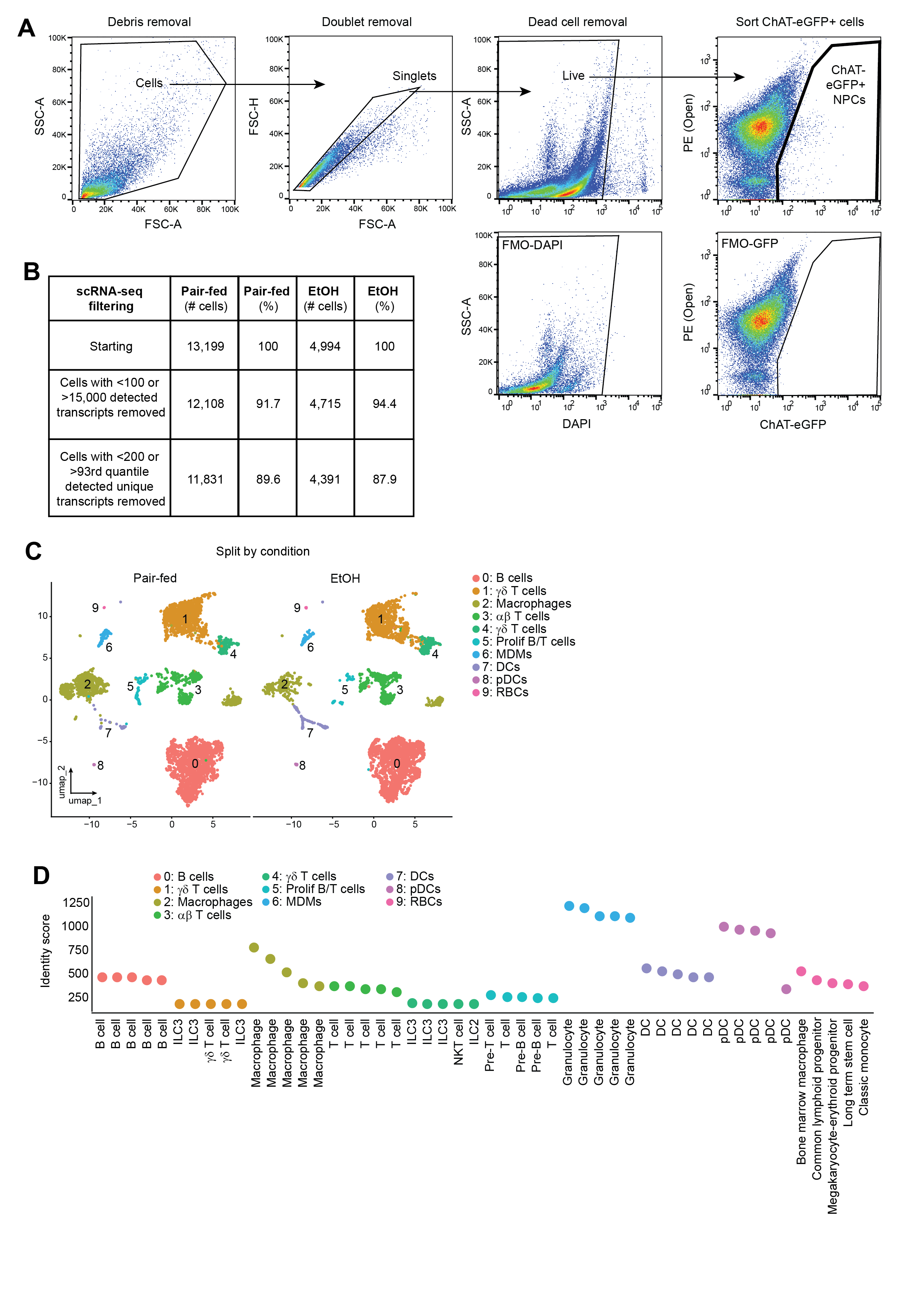

**Supplemental Figure 1. scRNA-seq of ChAT-eGFP+ NPCs.** (**A**) Gating strategy for FACS-based sorting of ChAT-eGFP+ NPCs from liver. Debris, doublets, and dead cells (DAPI+) were excluded, then ChAT-eGFP+ cells were gated on and sorted with reference to a fluorescence-minus-one (FMO) sample lacking GFP. (**B**) Attrition table showing the number and percentage of cells lost at each step of quality control filtering, resulting in the final number of valid cells used per condition (Pair-fed or Ethanol, EtOH). (**C**) UMAP plots of cell clusters split by (left) condition. (**D**) Cell type identity annotations determined by the cluster identity predictor tool (CIPR), drawing from ImmGen mouse reference database. MDMs: monocyte-derived macrophages; DCs: dendritic cells; pDCs: plasmacytoid dendritic cells; RBCs: red blood cells.

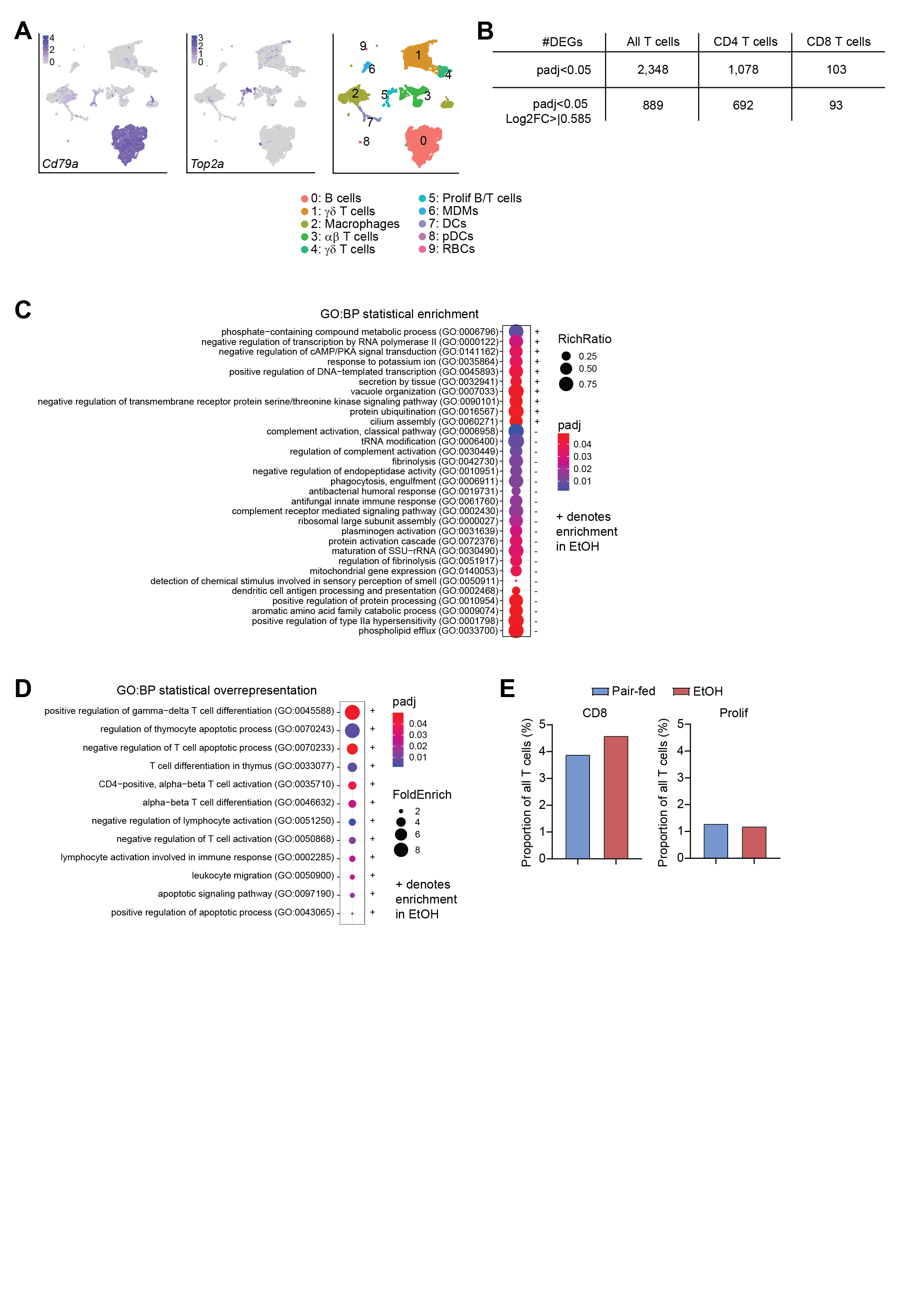

**Supplemental Figure 2. Impaired function of cholinergic T cells in liver stress.** (**A**) Feature plots showing *Cd79a* (B cells) and *Top2a* (proliferating cells) expression in all ChAT-eGFP+ cells. A UMAP of cluster names and numbers is shown on the right, from Supplemental Figure 1C. (**B**) Table showing the number of DEGs in all ChAT-eGFP+ T cells, CD4 T cells, or CD8 T cells, using two different thresholds (padj<0.05 or padj<0.05 with Log2FC>|0.585|). (**C**) Bubble plot of significantly enriched biological pathways derived from statistical enrichment analysis of ChAT-eGFP+ T cell DEGs (padj<0.05) between conditions. The Gene Ontology: Biological Processes (GO:BP) annotation set was used. The plus (+) symbol denotes terms that were enriched in the EtOH condition, and size and color of each bubble represent RichRatio and significance of each term, respectively. (**D**) Bubble plot of significantly deregulated biological pathways derived from statistical overrepresentation analysis of ChAT-eGFP+ T cell DEGs (padj<0.05 and Log2FC>|0.585|) between conditions. The Gene Ontology: Biological Processes (GO:BP) annotation set was used. The plus (+) symbol denotes terms that were enriched in the EtOH condition, and size and color of each bubble represent Fold Enrichment and significance of each term, respectively. (**E**) Relative proportions of CD8 and Prolif ChAT-eGFP+ T cell subsets in pair-fed versus EtOH conditions.

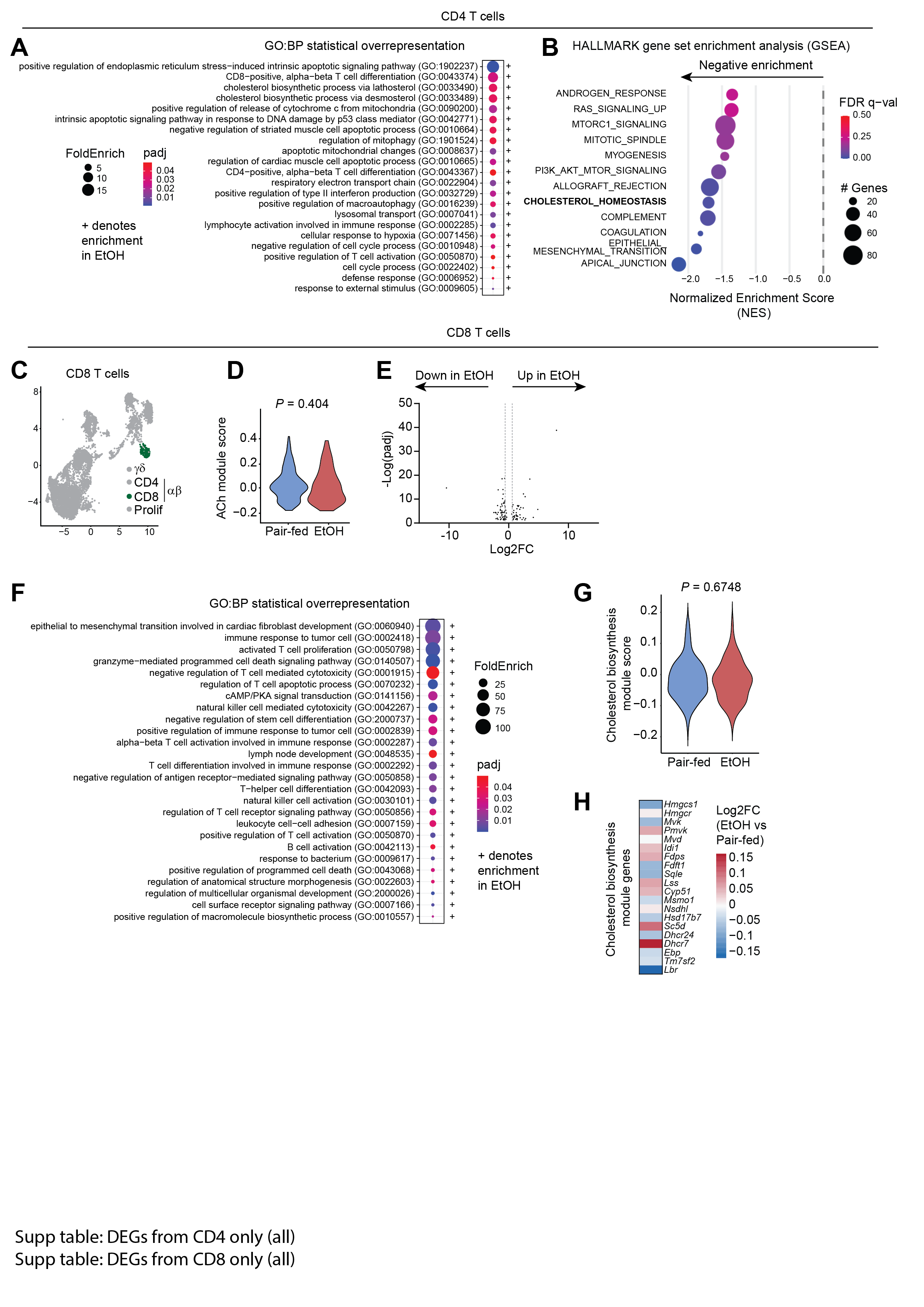

**Supplemental Figure 3. Cholinergic CD4 and CD8 T cell responses in liver stress.** (**A**) Bubble plot of significantly deregulated biological pathways derived from statistical overrepresentation analysis of CD4 T cell DEGs (padj<0.05 and Log2FC>|0.585|) between conditions. The Gene Ontology: Biological Processes (GO:BP) annotation set was used. The plus (+) symbol denotes terms that were enriched in the EtOH condition, and size and color of each bubble represent Fold Enrichment and significance of each term, respectively. (**B**) Gene set enrichment analysis (GSEA) of CD4 T cells between conditions, using mouse HALLMARK gene sets and all genes as input, regardless of significance. Normalized enrichment score (NES) was calculated, and terms with FDR q-val<0.25 are shown. Bubble color represents significance (FDR q-val) and size corresponds to the number of genes in each gene set that were found in the input gene list. (**C**) UMAP showing CD8 T cells colored dark green. (**D**) Composite expression score of an acetylcholine (Ach) gene module for CD8 T cells in the pair-fed or EtOH condition. (**E**) Volcano plot showing DEGs derived from differential expression analysis of CD8 T cells in the EtOH versus the pair-fed condition. Positive Log2FC indicates genes that were upregulated in EtOH. Vertical dashed lines intersect the x axis at Log2FC -0.585 and 0.585. (**F**) Bubble plot of significantly deregulated biological pathways derived from statistical overrepresentation analysis of CD8 T cell DEGs (padj<0.05 and Log2FC>|0.585|) between conditions. The Gene Ontology: Biological Processes (GO:BP) annotation set was used. The plus (+) symbol denotes terms that were enriched in the EtOH condition, and size and color of each bubble represent Fold Enrichment and significance of each term, respectively. (**G**) Composite expression score of a manually curated cholesterol biosynthesis gene module for CD8 T cells in the pair-fed or EtOH condition. (**H**) Heatmap showing fold enrichment (Log2FC) of each gene comprising the cholesterol biosynthesis module, for CD8 T cells from EtOH compared to pair-fed. For **D** and **G**, statistical testing was performed by Wilcoxon rank sum test with continuity correction and exact *P* values are shown above each graph.
